# The *Rpi-mcq1* resistance gene family recognizes *Avr2* of *Phytophthora infestans* but is distinct from *R2*

**DOI:** 10.1101/2020.10.08.331181

**Authors:** Carolina Aguilera-Galvez, Zhaohui Chu, Sumaiya Haque Omy, Doret Wouters, Eleanor M. Gilroy, Jack H. Vossen, Richard G.F Visser, Paul Birch, Jonathan D.G Jones, Vivianne G.A.A Vleeshouwers

## Abstract

Potato late blight, which is caused by the destructive oomycete pathogen *Phytophthora infestans*, is a major threat to global food security. Several nucleotide binding, leucine-rich repeat (NLR) *Resistance* to *P. infestans* (*Rpi*) genes have been introgressed into potato cultivars from wild *Solanum* species that are native to Mexico, but these were quickly defeated. Positional cloning in *Solanum mochiquense*, combined with allele mining in *Solanum huancabambense,* were used to identify a new family of *Rpi* genes from Peruvian *Solanum* species. *Rpi-mcq1*, *Rpi-hcb1.1* and *Rpi-hcb1.2* confer race-specific resistance to a panel of *P. infestans* isolates. Effector assays showed that the *Rpi-mcq1* family mediates a hypersensitive response upon recognition of the RXLR effector AVR2, which had previously been found to be exclusively recognized by the family of R2 resistance proteins. The *Rpi-mcq1* and *R2* genes are distinct and reside on chromosome IX and IV, respectively. This is the first report of two unrelated R protein families that recognize the same AVR protein. We anticipate that this likely is a consequence of a geographically separated dynamic co-evolution of *R* gene families of *Solanum* with an important effector gene of *P. infestans*.

**Author summary:** Potato is the largest non-grain staple crop and essential for food security world-wide. However, potato plants are continuously threatened by the notorious oomycete pathogen *Phytophthora infestans* that causes late blight. This devastating disease has led to the Irish famine more then 150 years ago, and is still a major threat for potato. Resistance against *P. infestans* can be found in wild relatives of potato, which carry resistance genes that belong to the nucleotide binding site-leucine-rich repeat (NLR) class. Known NLR proteins typically recognize a matching effector from *Phytophthora*, which leads to a hypersensitive resistance response (HR). For example, R2 from Mexican *Solanum* species recognizes AVR2 from *P. infestans*. So far, these *R* genes exclusively match to one *Avr* gene. Here, we identified a new class of NLR proteins that are different from R2, but also recognize the same effector AVR2. This new family of NLR occurs in South American *Solanum* species, and we anticipate that it is likely a product of a geographically separated co-evolution with AVR2. This is the first report of two unrelated R protein families that recognize the same AVR protein.

## Introduction

Potato (*Solanum tuberosum* L.) is the most important non-cereal crop directly consumed worldwide and plays a pivotal role in food security. To date, the major threat for potato production is the devastating late blight disease, which is caused by the Irish famine pathogen *Phytophthora infestans* [1]. Breeding for resistance to late blight began in the 1850s, when high levels of resistance were found in wild *Solanum demissum* that is native to the highlands of Toluca Valley in Mexico [2]. As a result of a tight co-evolution between wild *Solanum* and *P. infestans* in this center of genetic diversity, a wealth of resistance (*R*) genes have evolved in Mexican *Solanum* species [3–6]. Identified *R* genes include *R1*-*R11* from *S. demissum*, members from the *Rpi-blb1*, *Rpi-blb2* and *Rpi-blb3* family from *S*. *bulbocastanum* and *S. stoloniferum*, *Rpi-amr3* from *S. americanum*, and few more [7, 8]. Unfortunately, all *R* genes identified so far have failed to provide durable resistance to the ‘*R* gene destroyer’ *P. infestans* [1]. A second center of genetic diversity of *P. infestans* and tuber-bearing *Solanum* species occurs in South America [5]. More recently, new *R* genes from those regions have been isolated as well, such as the *Rpi-vnt1* family from *Solanum venturii* in Argentina and *Rpi-ber* family from Bolivian *S. chacoense*, *S. berthaultii* and *S. tarijense* [9, 10]. In addition to these, various other South American wild *Solanum* species are resistant to *P. infestans* and could provide useful *R* genes against late blight [11].

The oomycete pathogen *P. infestans* secretes an arsenal of effectors in order to manipulate host defense responses. Cytoplasmic effectors are typically members of the RXLR class, consisting of modular proteins with an N-terminal signal peptide, a conserved Arg-X-Leu-Arg (RXLR) motif required for translocation inside of the host cells and a highly polymorphic C-terminal domain [12]. *Avr2* of *P. infestans* is a well characterized RXLR effector and member of a highly diverse gene family. AVR2 targets the plant phosphatase BSL1 that is implicated in positive regulation of the brassinosteroid (BR) signaling. BR-signaling elevates expression of the transcription factor CHL1, which acts as a negative regulator of immunity. AVR2 was found to play an important role promoting *P. infestans* virulence and suppressing the effect of Pattern-Triggered immunity (PTI). Therefore, *Avr2* is considered an important effector gene and thus drives selection for *R* genes that recognize it [13, 14].

*R2* resistance genes which mediate recognition of AVR2 have been cloned from a major late blight resistance locus on the short arm of chromosome IV [15, 16]. Ten R2 homologs were identified from the Mexican wild *Solanum* species *S. bulbocastanum, S. demissum*, *S. edinense*, *S. hjertingii*, and *S. schenckii* [17, 18]. In addition to these, AVR2 was recently found to be also recognized by *Solanum mochiquense* (mcq) and *Solanum huancabambense* (hcb) that originate from Peru [18]. The activation of hypersensitive cell death responses upon delivery of AVR2 suggests that additional *R* genes are present in these *Solanum* species.

In this study, we focused on a new family of *Rpi* genes from the Peruvian *S. mochiquense* and *S. huancabambense*. *Rpi-mcq1, Rpi-hcb1.1* and *Rpi-hcb1.2* were cloned by map-based cloning in combination with a gene allele mining approach. The newly isolated *Rpi* genes encode NLR proteins that are sequence-unrelated with the *R2* family and confer resistance to *P. infestans* isolates. We show here that diversifying selection is acting in those genes, which may be a consequence of an independent molecular arms race between wild populations of *P. infestans* and *Solanum* species from South America.

## Materials and Methods

### Plant materials

The interspecific *Solanum mochiquense* BC1 mapping population was developed by crossing the *P. infestans*-resistant parent A988 (CGN18263) and susceptible parent A966 (CGN17731) [19]. The seeds were routinely treated with 1000 ppm gibberellic acid (GA3) for 24 hours to break dormancy and were sown on MS20 medium. Recombinants seedling plants were transferred to greenhouse and treated regularly with fungicides and pesticides to control other pests as described previously by Foster, Park (20).

### DNA isolation and sequencing

DNA isolation was performed using a Retch protocol [15]. BAC and cosmid clone DNA were isolated using the Qiagen Midi Prep Kit ® (Qiagen). BACs and cosmids ends, PCR products and shortgun clones were sequenced using the ABI PRISM BigDye® Terminator v3.1 Cycle Sequencing kit (Applied Biosystens) using manufacturer instructions.

### PCR-based marker development

The PCR-based markers used in this study are listed in SI Table **S1**. Cleaved amplified polymorphic sequence (CAPS) markers T0156 and TG328 were developed according with Smilde, Brigneti (19). TG591N and S1D11 were developed from restriction fragment length polymorphism (RFLP) markers annotated on Tomato-EXPEN map or Potato Maps [21]. Blast searches with the tomato gene sequences were performed on the *Solanum tuberosum* unigenes database at Sol genomics network [22] and the obtained Expressed Sequenced Taq (EST) homologs were used for CAPS markers development. U286446, U296361 and U272857 were developed from a released tomato BAC sequence (HBa_165P17) which overlaps with the *Rpi-mcq1* region. The CAPS marker 9C23R was developed from the BAC end sequence of the candidate 9C23 BAC clone. PCR products from resistance bulks and susceptible bulks were sequenced for each pair of primers. The identified SNPs were used to develop additional CAPS markers which were mapped on chromosome IX by using the 163 recombinants identified with T0156 and S1D11.

### BAC library construction and screening

A BAC library was constructed with pIndigoBAC-5 (*Hind III*-Cloning Ready) for a heterozygous resistant plant K182 carrying the *Rpi-mcq1* followed the instrument (Epicentre, WI, USA). The library is approximately 9× coverage with an average insert size of 85 kb. The BAC library was pooled for each 384-well plate and screened by PCR using the flanking and co-segregating makers. Positive clones were validated by singleton PCR.

### Allele mining

The *Rpi-mcq1*_*RGH* primers including the CACC site for Gateway® cloning purposes, were designed on *Rpi-mcq1* sequence (SI Table **S1**). Genomic DNA of the resistant genotype hcb353-8 was used as template in a PCR reaction (98C: 30‘′’, 25X [98C: 10‘′’, 64C: 30‘′’, 72C: 1‘30’′] 72C; 10‘′’) using Phusion® high-fidelity DNA polymerase (Thermo Fisher Scientific). Amplicons were separated on agarose gel and purified using the QIAquick Gel Extration Kit® (Qiagen). Purified products were cloned in pENTR/D-TOPO® (Invitrogen) using DH10B *E. coli* competent cells. The inserts of the entry clones were checked by PCR and Sanger sequencing. Unique sequences were transferred to the pK7WG2 destination vector [23] by an LR reaction using LR-clonase II® (Invitrogen) and plasmids were used for *A. tumefaciens* transformation strain AGL1.

### Binary vectors for*Agrobacterium tumefaciens*-mediated transient transformation

*Avr2*, *Avr3a*, *Rpi-hcb1.1* and *Rpi-hcb1.2* were cloned in the binary vector pK7WG2 [23]. *R2*, *Rpi-mcq1* and *R3a* were previously cloned in the binary vector pBINPLUS [24]. The NLR candidate genes cosA1, cosA2 and cosD5 were cloned in the binary cosmid vector pCLD04541 [25].

### Agroinfiltration

Agroinfiltration was performed as described by Domazakis, Lin (26). Briefly, young and fully expanded leaves of 4-5-week-old potato and *N. benthamiana* plants were used. A suspension of *A. tumefaciens* strain AGL1 containing the appropriate expression vectors at an OD_600_ of 0.2 for potato and at an OD_600_ of 0.5-1 for *N. benthamiana* were infiltrated in leaf panels. Three plants per genotype and three leaves per plant were used in two biological replicates. Local symptoms were assessed at 3-4 dpi. The percentage of cell death was quantified using scores of 0%, 25%, 50%, 75% and 100%, based on observation of the infiltrated area.

### Transient complementation

Agroinfiltration of *A. tumefaciens* strains carrying *Rpi-mcq1*, *Rpi-hcb1.1* or *Rpi-hcb1.2* were performed in four-week-old *N. benthamiana* plants according with method described by Domazakis, Lin (26). Two days after inoculation, infiltrated leaves were detached and inoculated with *P. infestans* isolates IPO-0 and IPO-C (S2 Table). Infection symptoms were scored between 4-8 dpi.

### Stable transformation

*A. tumefaciens*-mediated stable transformation was performed with potato cv. ‘Désirée’ and tomato cv. ‘Moneymaker’. The cosmid vector pCLD04541 harboring the cosmids A2, A1 and D5 under the control of their native promoters, were used in potato and tomato transformation to test the function of the NLR candidate genes. pK7WG2:*Rpi-hcb1.1* and pK7WG2:*Rpi-hcb1.2* under the control of 35S promoter were used in potato transformation to test the function of *Rpi-hcb* genes. The transformation was performed using routine transformation protocols [27]. Transformants were selected after growth in greenhouse conditions (18-22°C, 16 h of light and 8 h of dark).

### *Phytophthora infestans* isolates, culture conditions and inoculum preparation

The *P. infestans* isolates used in this study are listed in **S2** Table. Isolates were retrieved from our in-house stock collection. Isolates were grown in the dark at 15°C on solid rye sucrose medium prior to the disease test [28]. To isolate zoospores for plant inoculations, sporulation mycelium was flooded with cold water and incubated at 4°C for 1-3 hours.

### Disease test

Leaves from 6-8-week-old plants grown in greenhouse conditions (18-22°C, 16 h of light and 8 h of dark) were detached and placed in water-saturated oasis in trays. The leaves were spot-inoculated at the abaxial leaf side with 10μl droplets containing 5*10^4^ zoospores per ml (in tap water). After inoculation, the trays were incubated in a climate chamber at 15°C with a 16h photoperiod. Development of lesions and presence of sporulation was determined at 5-6 dpi. Disease index was estimated using a scale from 1 to 9, ranging from expanding lesions with massive sporulation (1 to 3, susceptible), sporulation no clearly visible (4), sporadic sporulation only visible under the microscope (5), lesion with a diameter size ≥ 10 mm (6), occurrence of hypersensitive response (HR) between 3 to 10 mm (7), less abundant sporulation and smaller lesions, occurrence of HR (8, resistant) and to no symptoms (9, fully resistant).

In all the cases, three to five leaves were inoculated per isolate, with 12 inoculation spots per leaf for potato (Désirée) and tomato (Moneymaker) transformants and 3 spots for *N. benthamiana* experiments. At least three independent experiments were performed.

### Phylogenetic and positive selection analysis

A UPGMA-based tree was generated with the full amino acid sequence of 16 proteins, including Rpi-mcq1 and R2 family members. Bootstrap value was set equal to 100. The obtained UPGMA-based tree was displayed as circular unrooted cladogram using Geneious®9.1.2.

A codon-based analysis was conducted using PAMLX1.3.1 package [29]. Maximum-likelihood codon substitution models M0, M1, M2, M7 and M8 were used for the analysis. Positively selected sites detected by, Models M2 and M8 were identified using Empirical Bayes Statistics [30].

## Results

### High resolution mapping and cloning of *Rpi-mcq1*

*Rpi-mcq1* (formerly named *Rpi-moc1*) was previously mapped to a region of 15.8 cM on chromosome IX, flanked by the CAPS marker T0156 and an AFLP marker [19]. In order to fine map *Rpi-mcq1*, a BC1 mapping population from *S. mochiquense* accessions A988 and A966 was developed. A recombinant screen on 2502 individuals from the mapping population using the flanking markers S1D11 and T0156 was performed. 163 recombinant individuals were identified and characterized for resistance to *P. infestans* isolates 90128 and EC1, resulting in resistant and susceptible bulks. Subsequently, five CAPS markers including TG328, U286446, U296361, TG591N and U272857 were screened on the recombinants (**S1** Table). *Rpi-mcq1* was mapped to a narrow region flanked by U286446 and S1D11, and co-segregates with TG591N and U282757 (**S1** Fig). U286446, TG591N and U282757 were tested on a BAC library, and two overlapping BAC clones 9C23 and 43B09 derived from the resistant haplotype were identified (Fig **1a**). A new CAPS marker, 9C23R, was obtained from the end sequence of BAC clone 9C23 and was screened, revealing segregation of *Rpi-mcq1* with six recombinants in the bulks. The marker 9C23R is located on the opposite orientation of *Rpi-mcq1* at a distance of 0.24 cM (Fig **1a**, **S1** Fig). *Rpi-mcq1* was fine-mapped on the two overlapping BACs, 9C23 and 43B09. Subsequently, the two BACs were sequenced by TIGR and as a result two contigs that shared 62,395 and 114,083 bp, respectively, were obtained. In total eight ORFs were predicted in the two contigs, including four *Tm2*^2^-like NLR candidate genes and four non-NLRs, including a putative NAD dependent epimerase, a RNA-directed DNA polymerase, a retransposon, and a protein with unknown function. Two NLR candidate genes, cosA2 and cosA1, have full length ORFs with ~80-85% identity to *Tm2*^2^ at the amino acid level. A third NLR candidate gene, cosD5, represents a partial NLR with a truncated Coil-Coil (CC) domain. The fourth NLR candidate gene, cosE7, contains an early stop codon and was omitted from further study.

**Fig 1.**
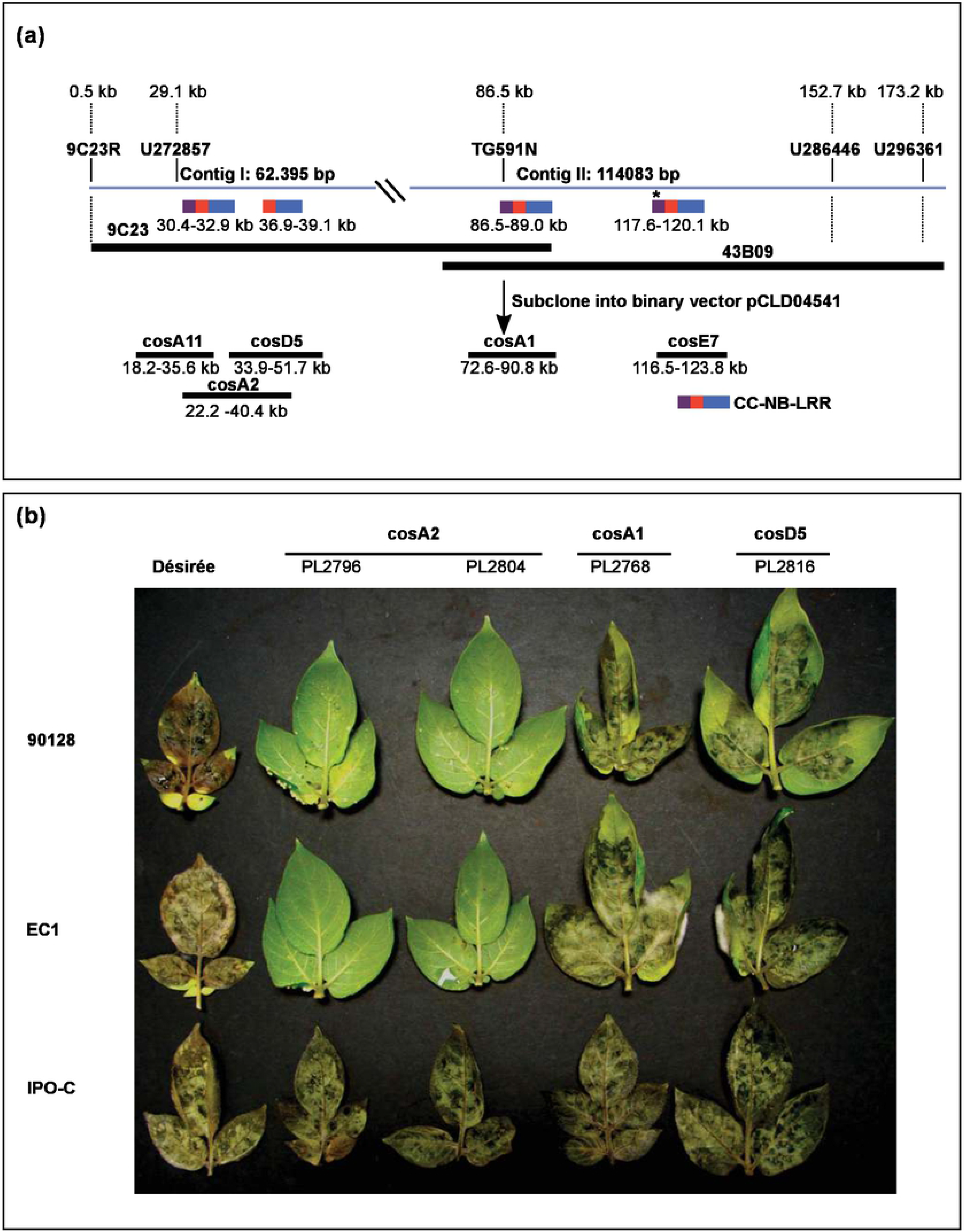
Physical map of the *S. mochiquense* genomic region containing *Rpi-mcq1* and genetic complementation. (a) Black rectangles represent bacterial artificial chromosome (BAC) clones (9C23 and 43B09) covering the region of *Rpi-mcq1* and cosmid clones carrying the NLR candidate genes. Blue lines indicate two contigs sequences from two BAC clones. Colored segments represent four NLR candidate genes, two of them encoded a complete CC-NB-LRR genes, represented as cosA2 and cosA1, respectively. Asterisk indicates an early stop codon in one NLR candidate gene. (b) Representative pictures of the disease symptoms of potato transgenic plants inoculated with *P. infestans* isolates 90128, EC1 and IPO-C, respectively. Pictures were taken at 6 dpi. PL2796 and PL2804 represent two transgenic lines obtained from cosA2 construct, PL2768 and PL2816 represent transgenic lines obtained with cosA1 and cosD5 constructs, respectively.

### *Rpi-mcq1* confers late blight resistance in potato and tomato

To determine whether the genes cosA2, cosA1 and cosD5 confer resistance to *P. infestans* in potato and tomato, stable potato and tomato transformants cv. ‘Désirée’ and cv. ‘Moneymaker’, respectively, were generated. In total, 22, 24, and 20 independent primary transformants were obtained in potato that express the cosA2, cosA1 and cosD5 genes, respectively, and 8, 7, and 10 independent tomato transgenic lines were obtained. Following transfer to the greenhouse, a detached leaf assay was performed on putative transformants and in the wild type ‘Désirée’. Detached leaves of the selected lines were spot-inoculated with the avirulent *P. infestans* isolates EC1 and 90128 (S2 Table). Macroscopic observations were carried out at 6 days post inoculation (dpi). On potato, 13 out of 22 putative transformants derived from cosA2 showed hypersensitive response (HR) and were resistant to the tested *P. infestans* isolates, whereas abundant sporulation was found in the 44 putative transformants derived from cosA1 and cosD5 and in the wild type ‘Désirée’ (Fig **1b**). Upon inoculation with the virulent *P. infestans* isolate IPO-C (S2 Table), all plants were infected (Fig **1b**). Consistent with these results, only the putative transformants derived from cosA2 showed race-specific resistance to EC1 and 90128 in tomato cv. ‘Moneymaker’ (S2 Fig). Altogether, the results indicate that cosA2 can complement the late blight susceptible phenotype in potato and tomato, and we designated it *Rpi-mcq1*.

### *Rpi-mcq1* homologs are detected in *Solanum huancabambense*

The wild *S. mochiquense* that carries *Rpi-mcq1* was previously shown to mount AVR2-specific cell death upon agroinfiltration [18]. In high-throughput effector screens in resistant wild *Solanum* germplasm, also *S. huancabambense* (hcb) was found to show AVR2-triggered cell death. To confirm the specificity of this response in independent experiments, *Avr2* was transiently expressed by agroinfiltration in hcb353-8 and in the wild type ‘Désirée’. Single infiltrations of empty vector pK7WG2 and co-agroinfiltrations of *R3a*/*Avr3a* were included as negative and positive controls, respectively. Specific cell death responses to AVR2 were evident in hcb353-8 and no cell death responses were identified on ‘Désirée’, suggesting the presence of an AVR2-recognizing R protein in hcb353-8 (**S3a** Fig).

To determine the resistance spectrum of hcb353-8, a detached leaf assay was performed with a panel of 12 *P*. *infestans* isolates. From this set, four and eight isolates are virulent or avirulent on *R2* plants, respectively (S2 Table). Among the twelve isolates, resistance phenotypes on hcb353-8 were fully correlating with *R2*-specific resistance (S2 Table). At 6 dpi, hcb353-8 showed a typical HR and was resistant to the avirulent isolate IPO-0, while large sporulating lesions were found in the cv. ‘Désirée’ (**S3b** Fig). These results indicate that a race-specific *Rpi* gene is present in hcb353-8, which exhibits similarities to AVR2-based resistance.

To investigate whether the *Rpi* gene in hcb353-8 is homologous to *Rpi-mcq1*, a genetic analysis was performed. The resistant hcb353-8 was crossed with the susceptible hcb354-2. Detached leaves of the parents and the 18 offspring genotypes were inoculated with *P. infestans* isolates 90128 and IPO-0. Macroscopic observations were carried out at 6 dpi. The susceptible hcb354-2 was infected by both isolates, whereas hcb353-8 showed HR and was resistant. No disease symptoms were found in ten genotypes of the progeny, whereas eight genotypes showed abundant sporulation and disease symptoms. This result confirms an expected 1:1 segregation ratio (x^2^= 0.11, P=0.74), suggesting the presence of a single dominant *Rpi* gene in hcb353-8 (**S3a** Table). Subsequently, to determine the genetic position of the *Rpi* in hcb353-8, a set of PCR markers that reside on LG IX of potato were tested on the F1 progeny (**S1** Table). The marker GP41 revealed polymorphism between the parents and progeny of the population and a complete co-segregation of marker patterns with *P. infestans* resistance was observed (**S3b** Table). Additionally, we agroinfiltrated the progeny plants with Avr2 and we found that the response to AVR2 segregates with resistance to *P. infestans* (**S3c** Table). In summary, the *Rpi* in hcb353-8 is located on LG IX and together with the AVR2-based race-specific resistance, we hypothesize that the *Rpi* gene in hcb353-8 is homologous to *Rpi-mcq1*. We followed a homology-based cloning approach to identify the *Rpi* gene. PCR with the conserved *Rpi-mcq1* primers (*Rpimcq1*_*RGH*) (**S1** Table) on genomic DNA of hcb353-8 plants yielded amplicons of 2577 bp, which is the range of the expected size of an *Rpi-mcq1* homologue. Sequence analysis of the amplicons revealed five additional *Rpi-mcq1* gene homologues (*RGH*) from hcb353-8. Amino acid similarities among these homologues vary between 73.3% to 86.6%, however, the similarity to members of the R2 family is much lower, namely between 42.4% to 45.6% (Fig **2**, **S4** Table). The phylogenetic relationship between *Rpi-mcq1*, *RGH* and the *R2* homolog *Rpi-blb3* was examined. A neighbor joining (NJ) tree grouped *Rpi-mcq1* and the *RGH* in a single clade, separate from *Rpi-blb3* (**S4** Fig). This result confirms that the *Rpi-blb3/R2* and *Rpi-mcq1* families are sequence unrelated and represent separate clades.

**Fig 2.**
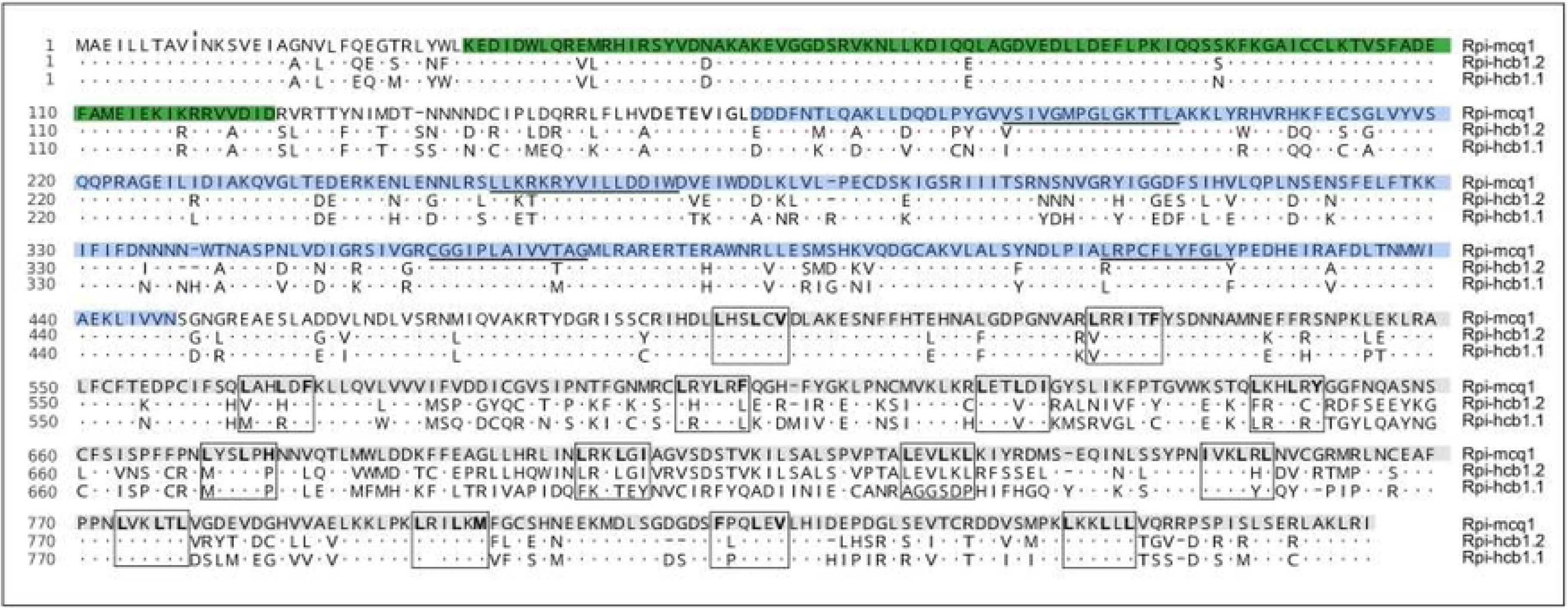
Protein alignment of the Rpi-mcq1 family. Multiple sequence alignment of Rpi-mcq1, Rpi-hcb1.1 and Rpi-hcb1.2, using the Rpi-mcq1 amino acid sequence as reference. The CC, NB-ARC and LRR domains are shaded in green, blue and grey, respectively. The characteristics conserved domains including the P-loop (Kinase 1a), Kinase 2, kinase 3 and the RNBS-D (resistance domain) in the NB-ARC are underlined. The 14 LRR repeats and the hydrophobic amino acids are represented with black rectangles and bold letters, respectively.

### *Rpi-hcb1.1* and *Rpi-hcb1.2* respond to AVR2 and confer resistance to *P. infestans*

To test the function of the *RGH*, the five identified homologues were cloned in binary vector pK7WG2. *Rpi-mcq1* was previously cloned in the binary vector pBINPLUS [18]. The constructs were transferred to *A. tumefaciens* strain AGL1. Co-agroinfiltrations were performed in potato cv. ‘Bintje’ leaves with *A. tumefaciens* expressing each *RGH* or *Rpi-mcq1* with *Avr2*. Single infiltrations of *Avr2*, *RGH* or *Rpi-mcq1* and empty vector were included as negative controls. Co-agroinfiltration of *R3a*/*Avr3a* was included as positive control. Two *RGH, i.e. RGH4* and *RGH5* induced specific cell death with AVR2 and we designated them *Rpi-hcb1.1* and *Rpi.hcb1.2,* respectively (Figs **2** **and** **3**).

**Fig 3.**
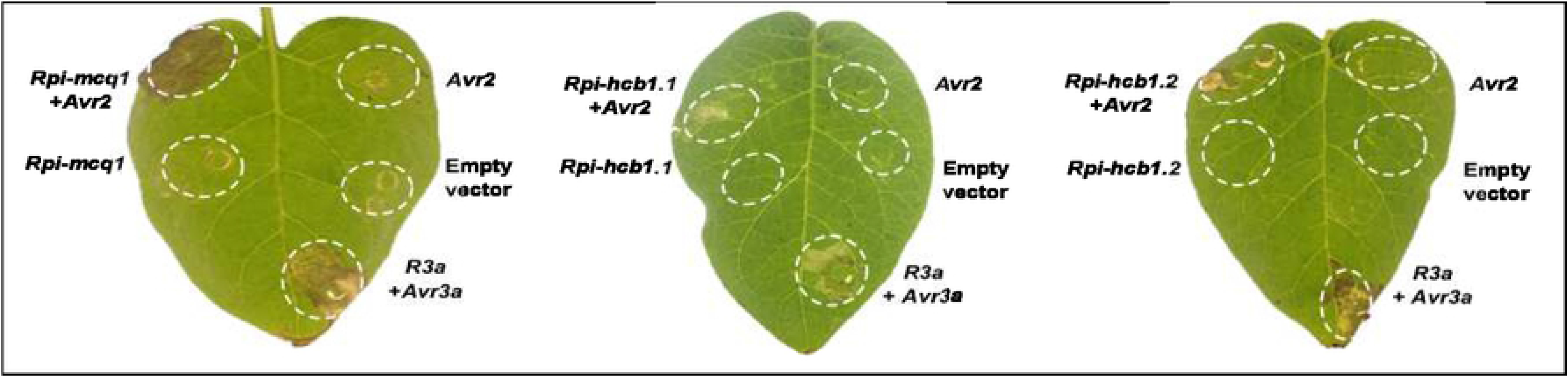
*Rpi-mcq1* homologues mediate response to AVR2. *Agrobacterium*-mediated expression of *Avr2* with *Rpi-mcq1*, *Rpi-hcb1.1* and *Rpi-hcb1.2* in potato cv. ‘Bintje’. Single infiltrations of each *R* gene alone, *Avr2* and empty vector were included as negative controls, and co-agroinfiltrations of *R3a*/*Avr3a* as positive control.

For assessing whether the *Rpi*-*hcb* genes confer resistance to *P. infestans*, a transient complementation assay was conducted. *Rpi-mcq1*, *Rpi-hcb1.1* and *Rpi-hcb1.2* were agroinfiltrated in *N. benthamiana* leaves. Additionally, agroinfiltrations with *Rpi-blb3*, empty vector, and infiltration medium (MMA) were used as a positive and negative controls, respectively. Two days later, zoospore suspensions of the avirulent IPO-0 and virulent IPO-C *P. infestans* isolates (**S2** Table) were spot-inoculated on the agroinfiltrated leaf panels [18]. After 8 days, *Rpi-*mcq1, *Rpi-hcb1.1*, *Rpi-hcb1.2,* and *Rpi-blb3-*treated leaf panels display a HR to IPO-0 (race 0), whereas large expanding necrotic lesions surrounded by sporulation zone were observed on leaves with the complex race IPO-C. Leaf panels agroinfiltrated with the empty vector and MMA medium show large sporulating lesions for both isolates (**S5** Fig). These data show that *Rpi-hcb1.1* and *Rpi-hcb1.2* confer a race-specific resistance to *P. infestans*, similar to *Rpi-mcq1-* and *Rpi-blb3-*mediated resistance.

To confirm the results with the transient complementation assay in potato, stable potato transformants cv. ‘Désirée’ were generated that express the *Rpi*-*hcb* genes under the control of the 35S constitutive promoter. 18 *Rpi-hcb1.1* and 24 *Rpi-hcb1.2* independent primary transformants were selected and cultured in greenhouse. Subsequently, they were tested for AVR2 response by agroinfiltration. Four and five independent transgenic lines expressing *Rpi-hcb1.1* or *Rpi-hcb1.2,* respectively, were selected. Detached leaves of the selected lines were inoculated with two avirulent *P. infestans* isolates PIC99183 and IPO-0 (**S2** Table). Stable transformed plants containing the *R* genes were resistant to both tested isolates. This result confirms that *Rpi-hcb1.1* as well as *Rpi-hcb1.2* confer resistance to *P. infestans* in potato (Fig **4**).

**Fig 4.**
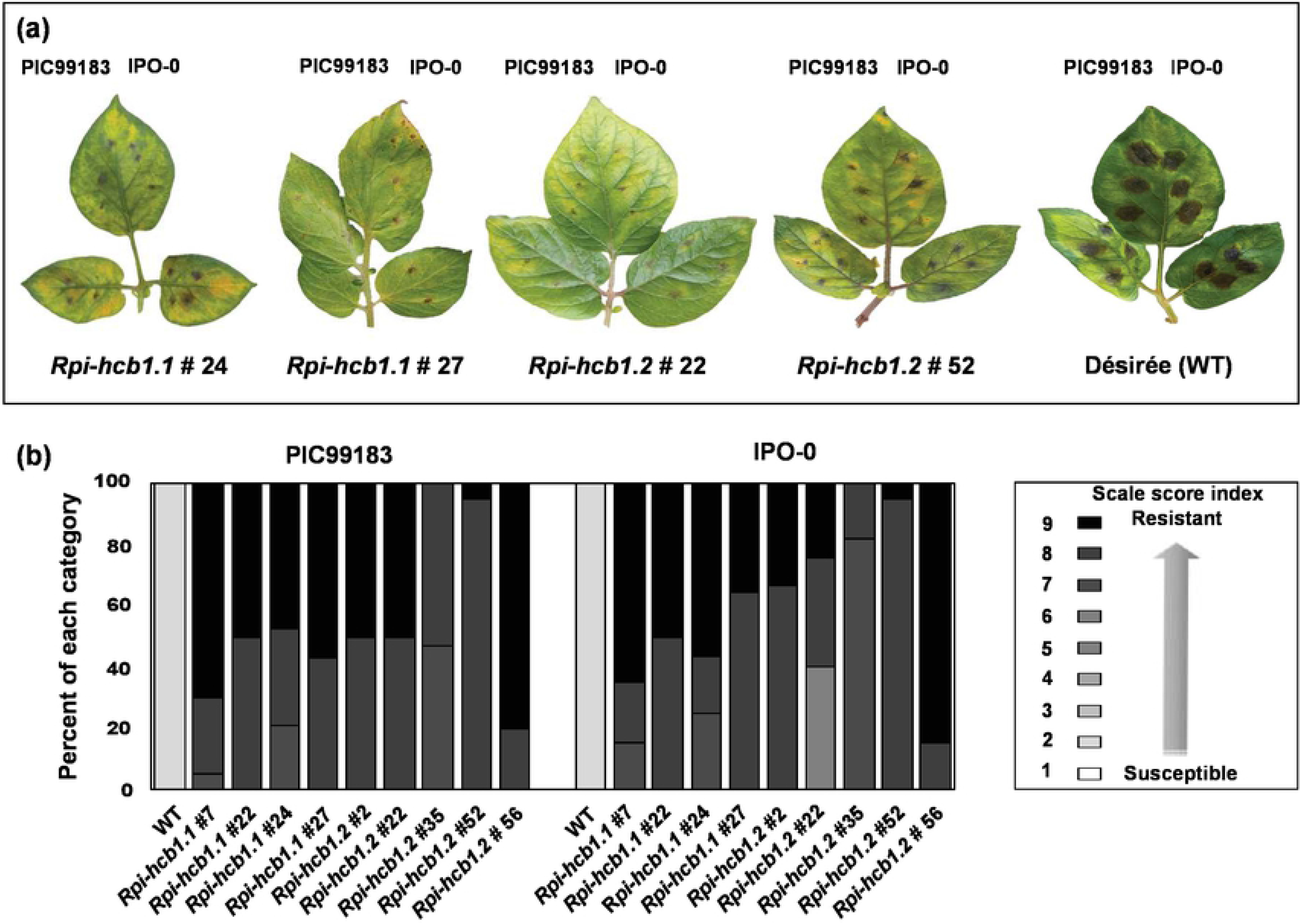
*Rpi-hcb1.1* and *Rpi-hcb1.2* confer late blight resistance. Four and five transgenic Désirée lines expressing *Rpi-hcb1.1* or *Rpi-hcb1.2,* respectively, were inoculated with *P. infestans* isolates PIC99183 and IPO-O. (a) Representative photographs are shown for Désirée expressing *Rpi-hcb1.1* lines 24 and 27 and *Rpi-hcb1.2* lines 22 and 52, respectively, at six dpi. Wild type (wt) ‘Désirée’ represents a susceptible control. (b) Disease symptoms on ‘Désirée’ (WT), Désirée-*Rpi-hcb1.1* lines 7, 22, 24 and 27 and Désirée-*Rpi-hcb1.2* lines 2, 22, 35, 52 and 56, are scored on a scale from 1 (susceptible) to 9 (resistant).

### *Rpi-mcq1* is under diversifying selection

The *Rpi-mcq1* family harbors all characteristics of NLR genes, and overall, the six *Rpi-mcq1* family members display considerable sequence conservation in the CC and NB-ARC domain. In contrast, the LRR domain, that is typically involved in effector recognition, contains multiple single nucleotide polymorphisms (SNPs). To get further insights on the region of *Rpi-mcq1* that is under diversifying selection, the PAML method was conducted [30]. Several positive selected amino acid residues were identified on the CC, the NB-ARC and the LRR domain (**S6** Fig). Model M2 identified 11 amino acid residues under positive selection, from which 10 amino acids were present in the LRR domain and 1 in the NB-ARC domain. The selection model M8 predicted the same 11 amino acids identified in M2 model and 6 additional amino acids. 1 out of 6 was present in CC domain, and the other amino acids were present in the LRR domain. This points to a strong diversifying selection on the LRR domain (**S6** Fig). This suggests that the polymorphisms could be driven by the evolution of differential specificities of AVR recognition that is typically located in the LRR domain [31].

## Discussion

Despite cloning of a number of *Rpi* genes in the past years [10], the fast-evolving *P. infestans* remains the most threatening pathogen of potato worldwide. To avoid losing the arms-race with this fast-evolving oomycete pathogen, it is necessary to intensify the search for new sources of resistance and extend the repertoire of available *Rpi* genes for breeding. The majority of introgressed resistance genes into potato cultivars, have been cloned from *Solanum* species native to Mexico, however, South America that is considered the second center of genetic diversity has been much less explored. In this manuscript, we report the cloning and functional characterization of a new family of late blight resistance genes from the Peruvian *S. mochiquense* and *S. huancabambense. Rpi-mcq1*, *Rpi-hcb1.1* and *Rpi-hcb1.2* confer race-specific resistance to *P. infestans* and belong to the *Rpi-mcq1* locus on linkage group (LG) IX [19]. The three *Rpi*-*mcq1* homologs share high levels of amino acid sequence identity, and we found that all three Rpi proteins mediate response to AVR2 of *P. infestans*.

Known *Avr* genes of *P. infestans* typically seem to have co-evolved with a single *R* locus in potato, such as *R3a* and *Avr3a* [32, 33]. However, AVR2 is the first example of an AVR protein that is recognized by two different, sequence-unrelated R protein families in tuber-bearing *Solanum* species. In Mexican *Solanum* species, AVR2 recognition is conferred by *R2* family on chromosome IV, whilst in Peruvian *Solanum* genotypes AVR2 response is mediated by *Rpi-mcq/hcb* family located on chromosome IX [18]. Since *Avr2* has a strong virulence function and is broadly spread among the pathogen strains, multiple *R* gene families seem to have evolved to recognize this effector [13, 14]. The *R* gene families display dissimilar sequences, the *Solanum* species occur in different phylogenetic clades and the plants occur in different geographic regions, indicating that they have evolved independently [18]. Our evolutionary analyses with Rpi-mcq/hcb sequences indicate that the LRR motif, which plays a pivotal role in effector recognition specificity, is shaped by a strong diversifying selection that could result in new specificities of effector recognition [17, 34]. Similar properties were found for the *R2* family [17]. These findings are in line with the concept of fast nucleotide evolution and sequence interchange between *R* gene homologs, as major mechanism shaping *R* gene diversity in plants, and especially targeted at the LRR [35, 36].

The recognition of an AVR protein by different R proteins has been reported for other plant-pathosystems, however, the relationship of those *R* genes has so far been unknown. For instance, *Avr3a/5* of *Phytophthora sojae* is recognized by the soybean resistance genes *Rps3* as well as *Rps5* from which the sequence is still unreported [37]. *Avr-Rmg7/8* of *Magnaporthe grisea* is recognized by the wheat resistance genes *Rmg7* and *Rmg8*, which are located in homeologous chromosomes 2B and 2A, respectively [38]. Furthermore, some rice *R* genes were recently found to interact with divergent *Magnaporthe oryzae* effectors via different binding surfaces [39, 40]. The effectors have evolved independently to unconventional *R* gene domains and target them with different binding-specificities [41]. These studies are providing more insights in the antagonistic interplay between pathogen and host that is driven by co-evolutionary forces targeted at *R* and *Avr* genes.

Pathogen populations in European countries, the USA and Canada have developed virulence to potato plants carrying the *S. demissum*-derived *R1*, *R3*, *R4*, *R7*, *R10* and *R11*. However, lower frequencies were noted for virulence on *R2,* and *R2* significantly delays the onset of epidemics in field trails [42, 43]. Since these characteristics heavily rely on the matching *Avr* gene, also *Rpi-mcq/hcb* are expected to display extended latency periods in the field and contribute to late blight resistance in practice.

Our improved understanding of the recognition of *Avr2* by *R2* and *Rpi-mcq/hcb* confirms that effectors are a powerful tool to accelerate the identification of *R* genes. This study is sharpening the concept that effector-based breeding should be complemented with additional molecular techniques, e.g molecular markers, to securely determine the identity of the matching *R* genes. Combination of *Rpi* genes from distinct loci but with similar race-specific resistance specificities, such as *R2* and *Rpi-mcq1,* could be more effective than using single *R* genes, however, a broader resistance spectrum is not expected based on AVR2 recognition alone. However, *Avr2* is member of a highly diverse gene family, and molecular studies on *Avr2* variants will reveal deeper insights underlying recognition specificities and pathogenicity in *P. infestans* populations. Such knowledge should contribute to wiser strategies for efficient deployment of *R* genes in potato.

## Acknowledgements

This work was supported by NWO-VIDI grant 12378 (V.G.A.A.V), COLCIENCIAS doctoral grant 617-2013 (C.A-G), The Veenhuizen Tulp Fund (C.A-G), COST action FA1208 (V.G.A.A.V, C.A-G, E.M.G, P.B., J.D.G.J). We thank Gert van Arkel for technical assistance, Isolde Pereira for plant maintenance, Geert Kessel, Francine Govers and David Cooke for providing *P. infestans* strains. Dionne Turnbull and Susan Breen are acknowledged for assistance in preliminary co-agroinfiltrations.

## Supporting information

**S1 Fig. High-resolution genetic linkage map of the *Rpi-mcq1* locus on chromosome IX.**

Genetic distances and marker names are indicated on the left and right of the map, respectively. *Rpi-mcq1* co-segregates with polymorphism makers TG591N and U272857, flanked by 9C23R and U286446 (or U296361), respectively.

**S2 Fig. Genetic complementation for late blight resistance on tomato with NLR candidate genes**. Representative pictures of the disease symptoms of PL3320, PL2741 and PL2759 tomato transgenic lines expressing the NLR candidate genes cosA2, cosA1 and cosD5, respectively. Detached leaves were inoculated with *P. infestans* isolates 90128 and EC1 and macroscopic observations for disease symptoms were carried out at 6 dpi.

**S3 Fig. *Solanum huancabambense* recognizes AVR2 and is resistant to *P. infestans***. (a) Agroinfiltration of *A. tumefaciens* carrying pK7WG2:*Avr2* and co-infiltrations of *R3a*/*Avr3a* on hcb353-8 and the wild type ‘Désirée. Single infiltrations of the pK7WG2 empty vector and co-infiltrations of *R3a*/*Avr3a* and/or *R2*/*Avr2* were included as negative and positive controls, respectively. (b) hcb353-8 shows no lesion development upon inoculation with *P. infestans* isolate IPO-O, whereas ‘Désirée’ (wt) shows large sporulating lesions at 6 dpi. Representative pictures are presented.

**S4 Fig. Phylogenetic relationship and sequence similarity of Rpimcq1 family.** The UPGMA-based tree illustrates the phylogenetic relationship at the amino acid level of Rpi-mcq1 and RGH cloned from hcb353-8 (blue) and the R2 family (red). Numbers on each node represent bootstrap values based on 100 replicates.

**S5 Fig. Genetic complementation of *Rpi-mcq1, Rpi-hcb1.1* and *Rpi-hcb1.2* in *N. benthamiana***. *Rpi-mcq1, Rpi-hcb1.1* and *Rpi-hcb1.2* were transiently expressed in *N. benthamiana* leaves, followed by inoculation with the *P. infestans* isolates IPO-0 and IPO-C. *Rpi-blb3* was included as a resistant control and infiltrations with medium (MMA) and with empty vector were used as negative controls. Race-specific resistance to IPO-O was observed with *Rpi-mcq1*, *Rpi-hcb1.1*, *Rpi-hcb1.2* and in the resistant control *Rpi-blb3*, whereas IPO-C caused expanding lesions.

**S6 Fig. Positive selection in **Rpi-mcq1** variants has mostly targeted the LRR domain**. (a) Graphical representation of posterior probability of diversifying selection based on model M8 at each site of Rpi-mcq1 variants. The * indicates a posterior probability higher than 99% of having *ω*>1. The x axis denotes codon position in the alignment of Rpi-mcq1 variants made from codeml and removing all the gaps. (b) * Log likelihood value. ** Likelihood radio test: = 2(lnl_alternative hypothesis_-lnl_null hypothesis_), with significance evaluated from distribution: df is degree of freedom and p is the probability. *** Bayes Empirical Bayer (BEB) analysis; amino acid sites, based on Rpi-hcb1.1 sequence, inferred to be under diversifying selection with probability >95%, and 99% in bold.

**S1 Table. Overview of CAPS markers and primers used in this study.** Primer sequence, annealing temperature and restriction enzyme are indicated. ^a^ Orientation of the primer: F, Forward; R, reverse. ^b^ Annealing temperature. ^C^ Restriction enzyme that reveals polymorphism between alleles linked to resistance or susceptibility.

**S2 Table. List of *P. infestans* isolates used in this study.**

The country, year, the genotype source of collection, the race, as well as the virulence (V) and avirulence (A) on *S. huancabambense* (hcb) 353-8 and *S. hjertjingii* (hjt) 349-3 plants are indicated. Hjt349-3 contains an *R2* homolog [18]

**S3 Table. Co-segregation for resistance, AVR2 response and genetic marker in hcb 7393 population.** (a) Resistance (R), susceptibility (S) or quantitative resistance (Q) against *P. infestans* isolates IPO-0 and 90128. (b) Presence (1) or absence (0) of the polymorphic band of the genetic marker GP41on chromosome IX. (c) Presence (+) or absence (−) of cell death response after agroinfiltration with *Avr2,* empty vector (negative control) and co-infiltration with *R3a/Avr3a* (positive control) at 4 dpi. n.d. not determined.

**S4 Table. Sequence similarities between Rpi-mcq1/hcb and R2 family.** Percentages of amino acid sequence similarities between ten R2 family members and five identified resistance gene homologues (RGH) of Rpi-mcq1 are presented.

